# Immunological responses to the relapsing fever spirochete *Borrelia turicatae* in infected Rhesus macaques: implications for pathogenesis and diagnosis

**DOI:** 10.1101/503615

**Authors:** Monica E. Embers, Aparna Krishnavajhala, Brittany Armstrong, Michael W. Curtis, Bapi Pahar, Hannah K. Wilder, Mike Allen, Paul Beare, Nicole R. Hasenkampf, Job E. Lopez

## Abstract

The global public health impact of relapsing fever (RF) spirochetosis is significant, as the pathogens exist on five of seven continents. The hallmark sign of infection is episodic fever and the greatest threat is to the unborn. With the goal of better understanding the specificity of B cell responses and the role of immune responses in pathogenicity, we infected Rhesus macaques with *Borrelia turicatae* (a new world RF spirochete species) by tick bite and monitored the immune responses generated in response to the pathogen. Specifically, we evaluated inflammatory mediator induction by the pathogen, host antibody responses to specific antigens, and peripheral lymphocyte population dynamics. Our results indicate that *B. turicatae* elicits from peripheral blood cells key inflammatory response mediators (IL-1β and TNF-α) which are associated with pre-term abortion. Moreover, a global decline in peripheral B cell populations was observed in all animals at 14 days post-infection. Serological responses were also evaluated to assess the antigenicity of three surface proteins, BipA, BrpA and Bta112. Interestingly, a distinction was observed between antibodies generated in non-human primates (NHPs) and mice. Our results provide support for the nonhuman primate model not only in studies of prenatal pathogenesis, but for diagnostic and vaccine antigen identification and testing.

## Introduction

Relapsing Fever (RF) spirochetosis is a neglected global disease. In parts of Africa, RF spirochetosis is a common bacterial infection [1], and the disease is a significant cause of hospital admissions and child mortality [2–6]. The causative agents are *Borrelia* species that are transmitted by the human body louse, or ixodid and argasid ticks (1–4). The manifestation of disease in humans includes recurrent febrile episodes, rigors, vomiting, severe headache, neurological symptoms, muscle and joint aches and tachycardia (1). Antibiotic treatment may result in the Jarisch-Herxheimer reaction, which is caused by a cytokine release leading to shock (5) and even death (6, 7). Mortality of tick-borne RF spirochetosis is 4–10% and is associated with the burden of spirochetes in the blood (8). RF borreliosis is particularly devastating on fetal and neonatal health (9, 10). For example, in Tanzania a perinatal mortality rate of 436/1000 was reported for *Borrelia duttonii* (11). The disease also has a severe impact in developing countries because of the nonspecific, malaria-like clinical manifestation of the disease. Importantly, with the geographic distribution of RF spirochetes largely overlapping with malaria (12) and studies indicating an often misdiagnosis (13, 14), the true morbidity of RF is underappreciated.

The reduction in spirochete levels and eventual clearance has been shown in animal models to be a direct result of the antibody response, especially IgM and IgG3 isotypes (15, 16). The clearance by lymphocytic response was established by Newman and Johnson (17), who showed not only the importance of the B cell response, but that of a T-independent B cell response. Subsequent studies have demonstrated neutralization (18) and a directly bactericidal (19) role of serum IgM in controlling relapsing fever spirochetemia. The contribution of B cell subsets to RF pathogen control has been further delineated in mice (16, 20, 21).

Rodent models of RF have contributed immensely to the understanding of infectivity, host-pathogen interactions and immune responses to infection [31–36]. For example, transmission studies in *Borrelia turicatae* demonstrated that RF spirochetes enter the host within seconds of tick bite (22), indicating the importance of preventing early mammalian infection. Moreover, vaccination of mice with the *Borrelia hermsii* variable tick protein (Vtp) has guided vaccine strategies. Vtp is produced in the salivary glands of *Ornithodoros hermsi* and subsequently down-regulated once the pathogens are detectable in murine blood (23). Vaccination studies with Vtp indicated that RF spirochete surface proteins produced in the tick salivary glands could be ideal immunological targets to prevent the establishment of infection (24).

Mice are natural reservoir hosts and may have limitations as models for testing intervention and therapeutic strategies. Thermoregulation in mice varies, and they are a limited model to further understand the Jarish-Herxheimer reaction. Mammals have evolved unique thermoregulatory mechanisms in defense against pathogens, with rodents typically remaining afebrile or decreasing body temperatures in response to bacterial challenge and endotoxin administration (25–29). Therefore, mice may not be ideal for the evaluation of vaccine candidates and therapeutics that prevent the clinical sign of fever, which is a hallmark feature of RF.

Non-human primates (NHP) infected with RF spirochetes accurately mimic human disease. A 1938 report published by Dr. Edward Francis showed that NHPs infected with *B. turicatae* by tick bite exhibited morbidity and mortality commonly observed with human disease (30). We have also demonstrated human-like illness with this model. Four rhesus macaques were infected with *B. turicatae* by tick transmission, and radio telemetry was used to quantify the intricacies of infection (31). Multiple febrile episodes, high spirochete densities in blood, and disruption of cardiac function were observed.

In this current report, we further characterized the immune responses of NHPs that were infected with *B. turicatae* by tick bite (31). We originally hypothesized that *B. turicatae* would induce a TH2 type immune response, with concomitant induction of B cell proliferation and antibody production. Rather, we found that in peripheral blood cells, *B. turicatae* induced TH1 type cytokines (IL-1β and TNF-α) and significant declines in B cell populations were observed soon after infection. Changes in peripheral blood lymphocyte subsets, immune mediator production by stimulated PBMCs, and antibody responses reflect a distinct response to RF *Borrelia* in NHPs. We evaluated antibody responses to a known conserved surface protein, the *Borrelia* immunogenic protein A (BipA) (33, 34), and two newly identified surface proteins, Bta112 and the *Borrelia* repeat protein A (BrpA). Bta112 and BrpA are up-regulated in the tick and were evaluated to determine their antigenicity once *B. turicatae* enters the mammalian host. Our results demonstrate differences in the host antibody specificity between mice and NHPs infected with *B. turicatae*, and further indicate the significance of macaques as a model that most accurately represents human RF borreliosis.

## Results

### Co-culture of macaque PBMCs with *B. turicatae* elicits inflammatory response mediators

Given the high numbers of RF spirochetes that are observed in the blood during febrile episodes, we sought to measure immune mediators produced by PBMCs in response to stimulation with borreliae. In the analysis of 23 cytokines produced in reponse to stimulation with *B. turicatae*, both commonalities and differences with *B. burgdorferi* (the Lyme disease causing agent) were observed. While both borrelial pathogens elicited TNF-α, IL-10, G-CSF and IL-12/23p40, *B. turicatae* induced a statistically significant higher level of IL-1β and soluble CD40 ligand (sCD40L) compared to *B. burgdorferi* (Figure 1A and 1D, respectively). We tested stimulation of PBMCs derived from a naïve, uninfected monkey (JD03) in addition to PBMCs derived from infected animals. To preserve the viability of the spirochetes and retain soluble factors, the stimulations were performed with spirochetes in their own growth media (shown as Bt media and Bb media). However, components of the media also had a moderate stimulatory effect for some cytokines/chemokines. Figure 1A shows IL-1β responses of naïve macaque PBMCs stimulated with borreliae, indicating a significant induction of this inflammatory cytokine specifically by *B. turicatae*. For IL-1β, significance differences were observed at the 12-hour time point when comparing *B. turicatae* (Bt) to BSK (p=0.0231) and *B. burgdorferi* (Bb) to BSK (p=0.001). At 24 hours, significant differences in these two groups were observed as well (Bt vs. BSK, p<0.0001; Bb vs. BSK, p=0.0047). In Figure 1B, the effect on TNF-α production indicates that both *Borrelia* species induce production of this inflammatory cytokine by PBMCs. At 12 hours, significance was observed when comparing Bt vs. BSK (p=0.0007), Bb vs.BSK (p<0.0001). No difference was observed between Bt media and BSK, yet the quantity of TNF-α induced by Bt over that of Bt media was significant (p=0.0013), indicating that soluble factors do not drive the induction of TNF-α by B. *turicatae*. At 24 hours, each of these differences remained significant (Bt vs. BSK, p=0.0002; Bb vs. BSK, p=0.0003; Bt media vs. Bt). Figure 1 also shows the specific differences in the induction of G-CSF (C), sCD40 (D) and IL-12/23 (E) from naïve PBMCs stimulated with *borreliae*. Significant changes in G-CSF production by PBMCs stimulated with *B. turicatae* were only observed at the 24 hour time point. Here, stimulation with Bt vs. BSK was significant (p=0.0005), as was stimulation with Bb vs. BSK (p=0.0002). For the soluble CD40 ligand, (sCD40l), significant differences were observed only at the 12 hour time point, with stimulation of Bt compared to Bt media demonstrating significance (p=0.0085), along with Bt vs. BSK (p=0.0027) and Bt vs. BSK (p=0.0047). For IL-12/23, significant differences were seen at both 12 and 24 hour time points comparing Bt vs. BSK (12 h: p=0.0007; 24h: p=0.0013) and Bb vs. BSK (12h: p=0.0033; 24 h: p=0.0343). Figure 2 shows IL-1β responses among infected monkeys. Stimulation of day 14 p.i. PBMCs with Bt vs. Bt media alone resulted in a significant increase in this inflammatory mediator for all three monkeys that were infected. A two to three fold increase in quantity of IL-1β produced in response to *B. turicatae* compared to media alone indicates the specific effect of the pathogen. Specifically, significant differences were observed at the 12 and 24 h timepoints for JB60 (12 h: p=0.0070; 24 h: p=0.0010), JB23 (12 h: p=0.0022; 24 h: p=0.0005), and IN57 (12 h: p=0.0008; 24 h: p=0.0005) when Bt vs. Bt media were compared.

**Figure 1.**
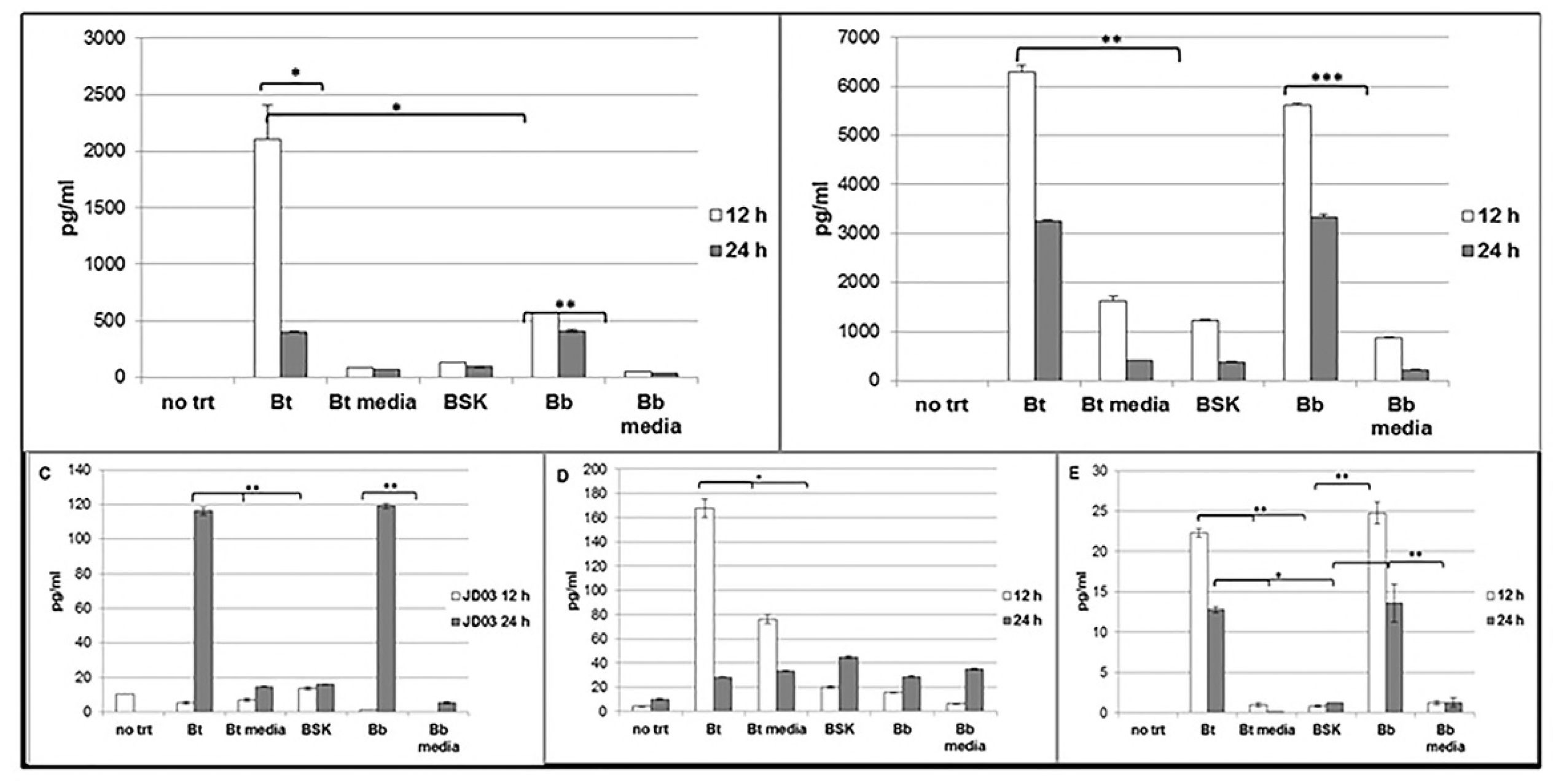
IL-1β (A),TNF-α (B), G-CSF (C), sCD40 (D) and IL-12/23 (E) responses of naïve macaque PBMCs stimulated with Borreliae. PBMCs obtained at day 0 from animal JD03 were stimulated with *B. turicatae* (Bt), *B. burgdorferi* (Bb), filtered BSK-H medium derived from *B. turicatae* or *B. burgdorferi* cultures (Btmedia; Bbmedia), uninoculated BSK-H medium (BSK), or left untreated (no trt). Supernatants were collected at 12 and 24 hours, for measurement of inflammatory mediators by a NHP-specific 23-plex cytokine bead assay.

**Figure 2.**
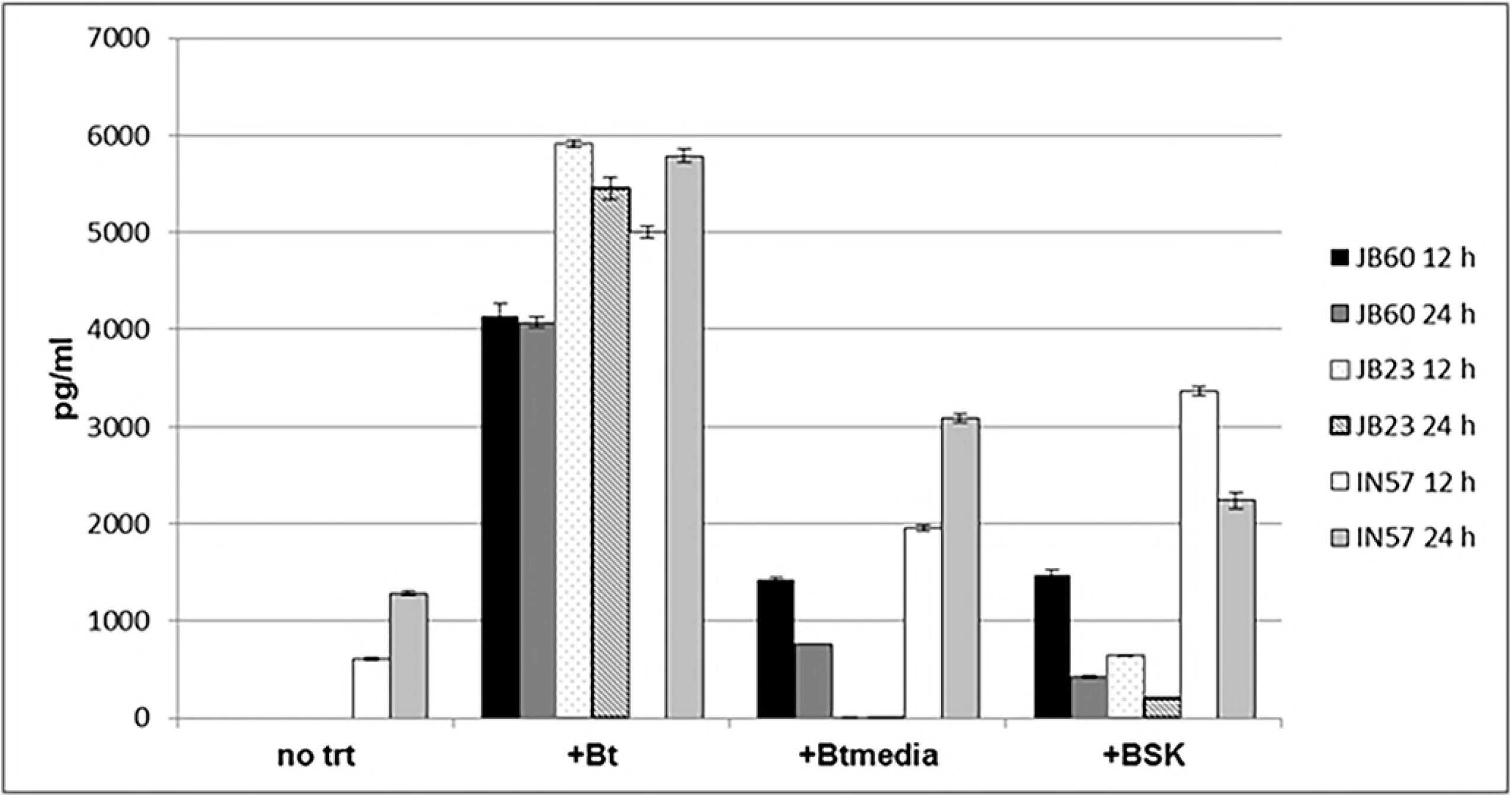
IL-1β production by the PBMCs of *B. turicatae-infected* macaques. Cells isolated from blood on day 14 post-tick feeding were incubated with *B. turicatae* at a 10:1 ratio of spirochetes to cells (+Bt), untreated (no trt), incubated with BSK-H medium (+BSK), or incubated with filtered mBSK medium derived from *B. turicatae* cultures (+Btmedia). Supernatants were collected at 12 and 24 hours, for measurement of inflammatory mediators by a NHP-specific 23-plex cytokine bead assay.

Immune regulatory molecules in serum were also quantified. We collected blood droplets between days 0–14, but serum was collected beginning on day 14, as our intent was to evaluate antibody responses. Therefore, we used available sera to test in the 23-plex cytokine magnetic bead panel. The immune mediators that were elevated in serum at various time points included IL-10, sCD40L, IL-8 and MCP-1 (supplemental Figure S2). Both IL-10 and MCP-1 were elevated at the earlier time points (14 and 28 days), but declined by 6 weeks post-infection. In contrast, sCD40L and IL-8 appeared to be elevated throughout the infection period in monkeys inoculated with *B. turicatae* (IN57, JB60 and JB23) compared to the animals fed upon by uninfected ticks (JD03).

### Characteristic peripheral B cell depletion during acute infection

T and B-cell phenotypic analyses were performed with PBMCs at days 0, 14, and 70 post-infection time points from all 4 NHPs. With respect to the T-cell phenotype, only general CD4 and CD8 T-cell phenotypes were measured in all three time points after infection. Only one animal (IN57) had reduced CD3+ populations, detected at day 14 post-infection (Table S1 and Figure 3) and the percentages of all four subsets of CD3 cells (CD4+CD8-, CD4+CD8+, CD4-CD8+ and CD4-CD8-) were reduced. The other 3 animals showed a moderate increase in peripheral T cells at 2 weeks p.i. that declined by day 70.

**Figure 3.**
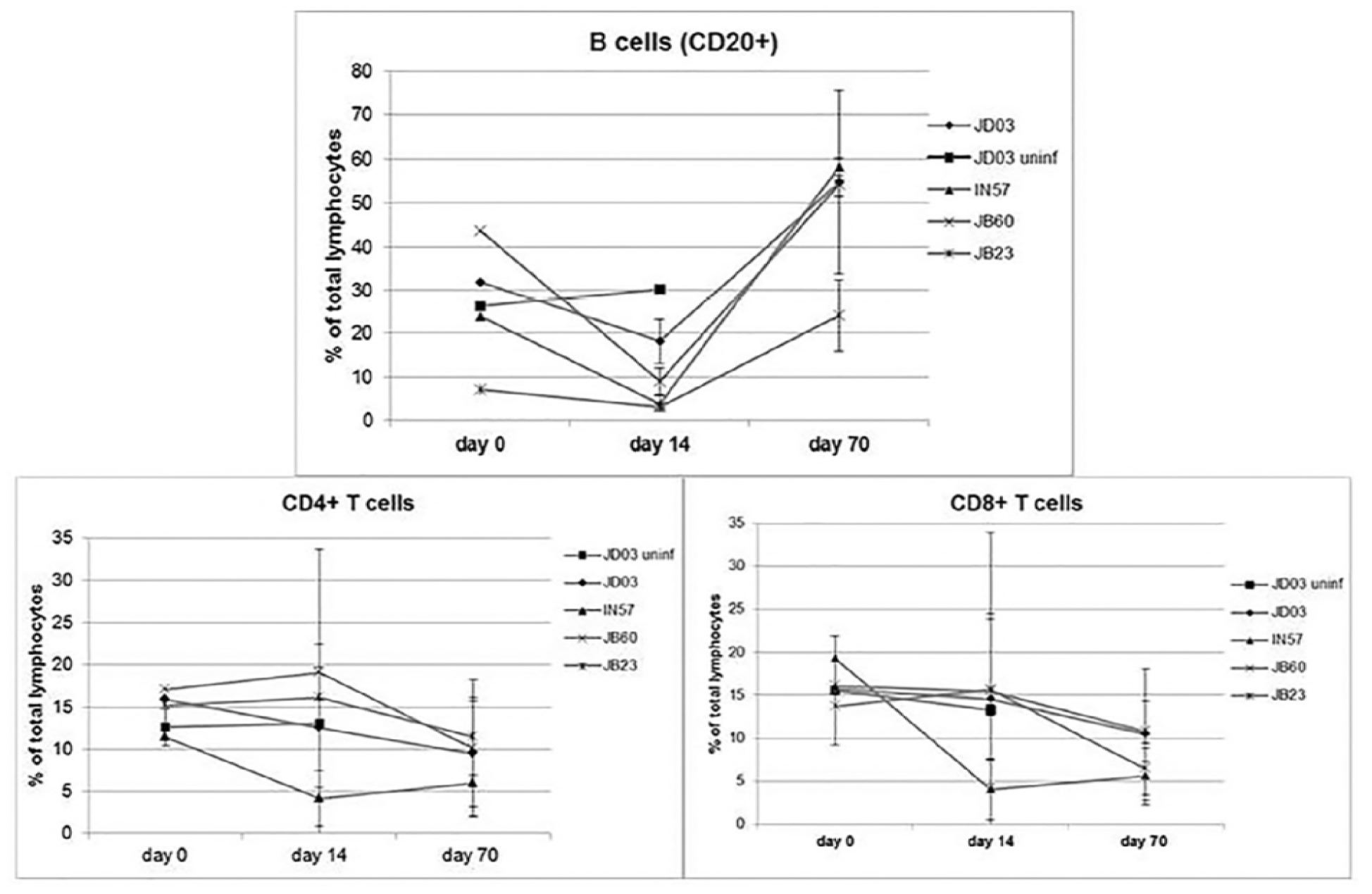
Frequency of B and T cells in the peripheral blood after infection with *B. turicatae*. PBMCs were subjected to flow cytometry to detect the relative percentages of B cells (CD20+) and CD4+/CD8+ T cell subsets. Each staining experiment was performed twice and the standard deviation is indicated with error bars.

B-cell subsets were distinguished by a panel of markers that included CD5, CD20, CD21, CD138, IgM, IgD, CD27, and CD38. Notable reductions in the percentages of B cells were observed in the serum 14 days after infection, suggesting an infection-induced B cell depletion (Table 1, Figure S1). As shown in Table 1, the B-cell depletion was due to the loss of CD5 (B-1a cells, marker of naïve or immature B-cells (35–37)), CD21 (marker of B cell differentiation and maturation (38–40)), CD86 (activation marker (41, 42)), and CD138 (plasmablasts (37)). In a subsequent staining, we looked at IgM+ B cells, switched memory (CD27+IgD-), non-switched memory (CD27+IgD+), naïve (CD27-IgD+), double negative (CD27-IgD-), and CD27^high^CD38^high^ plasmablasts. A precipitous and global decline in peripheral B cell populations was observed in all animals at day 14 p.i. (Table 1 and Figure 3). The B cell percentages and different B-cell subsets returned to near pre-infection levels at day 70 post-infection in all animals. The percent drop in total B cell frequency between day 0 and day 14 was significant for all monkeys. Specifically, the CD20+ lymphocytes decreased by 43% for JD03, 84% for IN57, 80% for JB60 and 56% for JB23.

### Evaluation of Bta112 between strains of *B. turicatae*

Bta112 was further evaluated as an antigen because computational analyses suggested the protein was exposed on the surface of RF spirochetes. The PROSITE InterPro database identified a predicted lipid attachment site at the N-terminus of the protein (Figure S3). PSIPRED and the Phobius prediction server suggested that the Bta112 was rich with alpha helices and the C-terminus of the protein was soluble and positioned toward the extracellular environment, respectively. Sequence analysis of Bta112 between *B. turicatae* 91E135, FCB, TCB1, TCB2, and 99PE-1807, indicated the presence of an intact gene that coded for a protein that was nearly identical in all *B. turicatae* isolates evaluated (Figure S3). Given the presence of Bta112 in multiple *B. turicatae* isolates, we evaluated the protein further.

### Expression of recombinant Bta112, temperature-mediated production, and surface localization of the native protein

To evaluate serological responses to *B. turicatae* Bta112, the gene was expressed as a recombinant fusion protein. *bta112* was overexpressed in BL21 Star (DE3) cells (Figure 4A), and rabbit immune serum was generated against the recombinant protein. Since native *bta112* is up-regulated by *B. turicatae* during culture at 22°C relative to 35°C (43), spirochetes grown at both temperatures were evaluated to assess temperature-mediated protein production. Optical density analysis of immunoblots probed with the rabbit serum sample generated against rBta112 indicated 3.2-fold increase of the protein in *B. turicatae* grown at 22°C versus 35°C (Figure 4B, top panel). The rabbit’s pre-immunization serum sample was used as a negative control (Figure 4B, middle panel). Moreover, a serum sample generated against *B. turicatae* FlaB was used as a control to indicate similar protein loads were electrophoresed in the immunoblotting assays (Figure 4B, lower panel).

**Figure 4.**
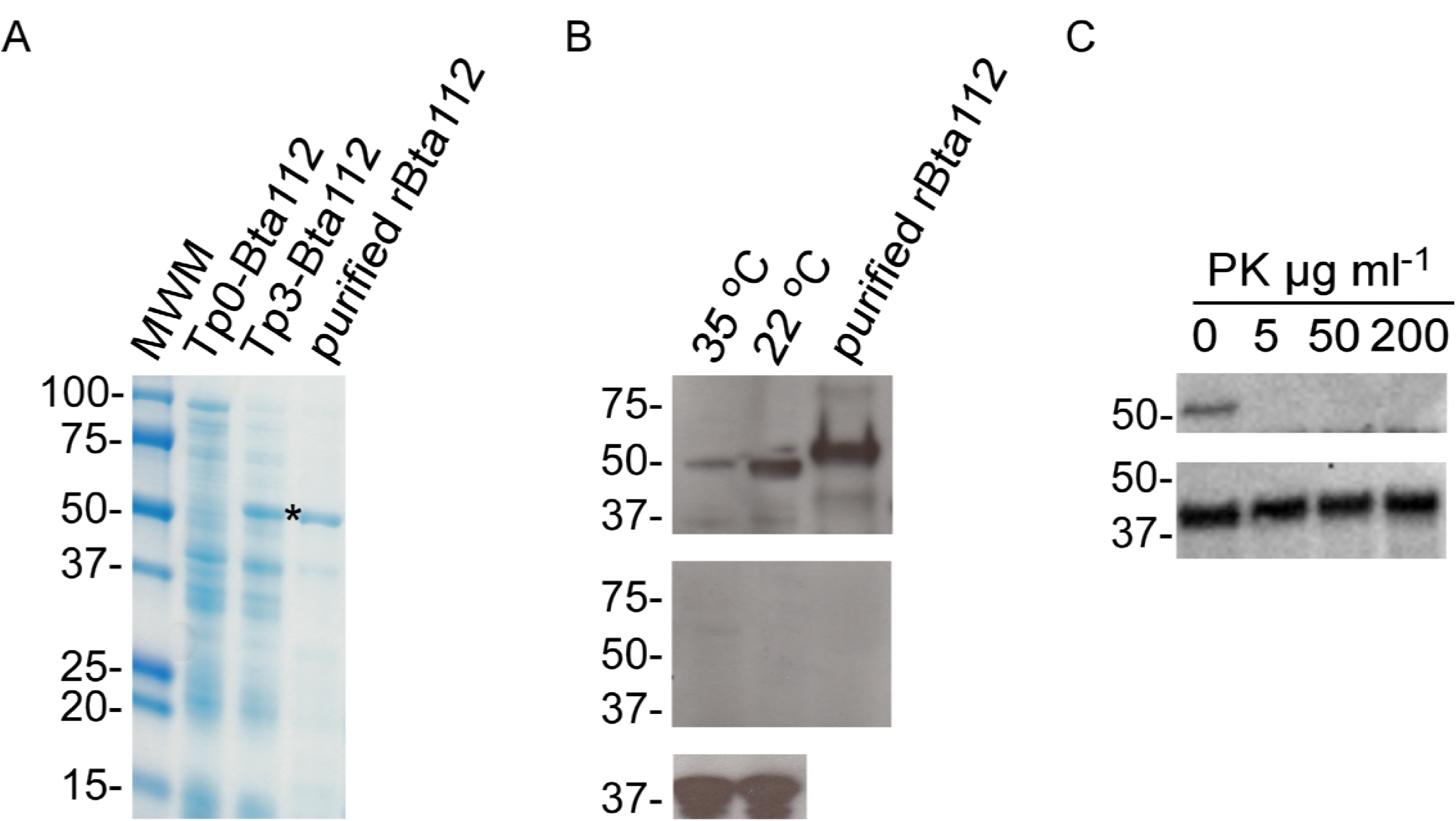
Expression of *bta112* as a recombinant protein, temperature mediated protein production and surface localization of Bta112. Bta112 was produced in *E. coli* and purified (A). *E. coli* samples were taken prior to induction (Tp0), three hours after induction (Tp3), and the purified protein. Immunoblots using rabbit serum samples generated against rBta112 indicated the protein’s increased production at 22°C compared to 35°C (B, upper panel). Preimmune serum samples (B, middle panel) and serum samples generated to the flagellin (FlaB) protein (B, lower panel) are shown. Immunoblotting was also performed to evaluate the surface localization of Bta112. Proteinase K (PK) was used at concentrations of 5 to 200 μg per ml (C). Membranes were probed with anti-Bta112 serum samples (C, upper panel), and anti-FlaB (C, lower panel) as a control to indicate equal protein loads in the gels (73).. Molecular weight markers (MWM) are show on the left of gel (A), and molecular masses are indicated on the left of each immunoblot.

Performing proteinase K and immunoblotting assays with *B. turicatae* grown at 35°C indicated that the Bta112 was surface localized (Figure 4C). Bta112 was degraded following incubation with increasing concentrations (5, 50, and 200 μg per ml) of proteinase K for 15 minutes (Figure 4C, upper panel). The relative density of the periplasmic protein FlaB in 5 μg per ml of proteinase K compared to the 0 μg per ml of proteinase K control was 101%. The relative density of FlaB in 50 and 200 μg per ml of proteinase K was 93% and 90% respectively, indicating that the spirochetes’ membranes remained intact (Figure 4C, lower panel). Collectively, these results supported that Bta112 was surface localized and the protein’s production was elevated at 22°C. Given these findings, the antigenicity of rBta112 was assessed.

### Serological responses to *B. turicatae* surface proteins

Given variations in humoral responses between mammalian species (44, 45), we compared the antigenicity of *B. turicatae* rBta112, rBrpA, and rBipA using serum samples from NHPs and mice that were infected by tick bite. Immunoblotting indicated varying serological responses between NHPs and mice to the recombinant proteins (Figure 5 A-F). All four NHPs produced antibodies that bound to *B. turicatae* protein lysates, rBta112, and rBipA, while serological reactivity to rBrpA was only detected in JB60 (Figure 5B). An immunoblot from two mice represented the eight remaining animals (Figure 5E and F). All the mice seroconverted to rBipA, two of eight animals seroconverted to rBta112, while none of the animals seroconverted to rBrpA. Probing immunoblots with a monoclonal antibody for the six histidine epitope was used to as a control for the expected molecular weight of each protein (Figure 5G). These findings suggested varying serological responses to RF spirochete antigens between NHPs and mice.

**Figure 5.**
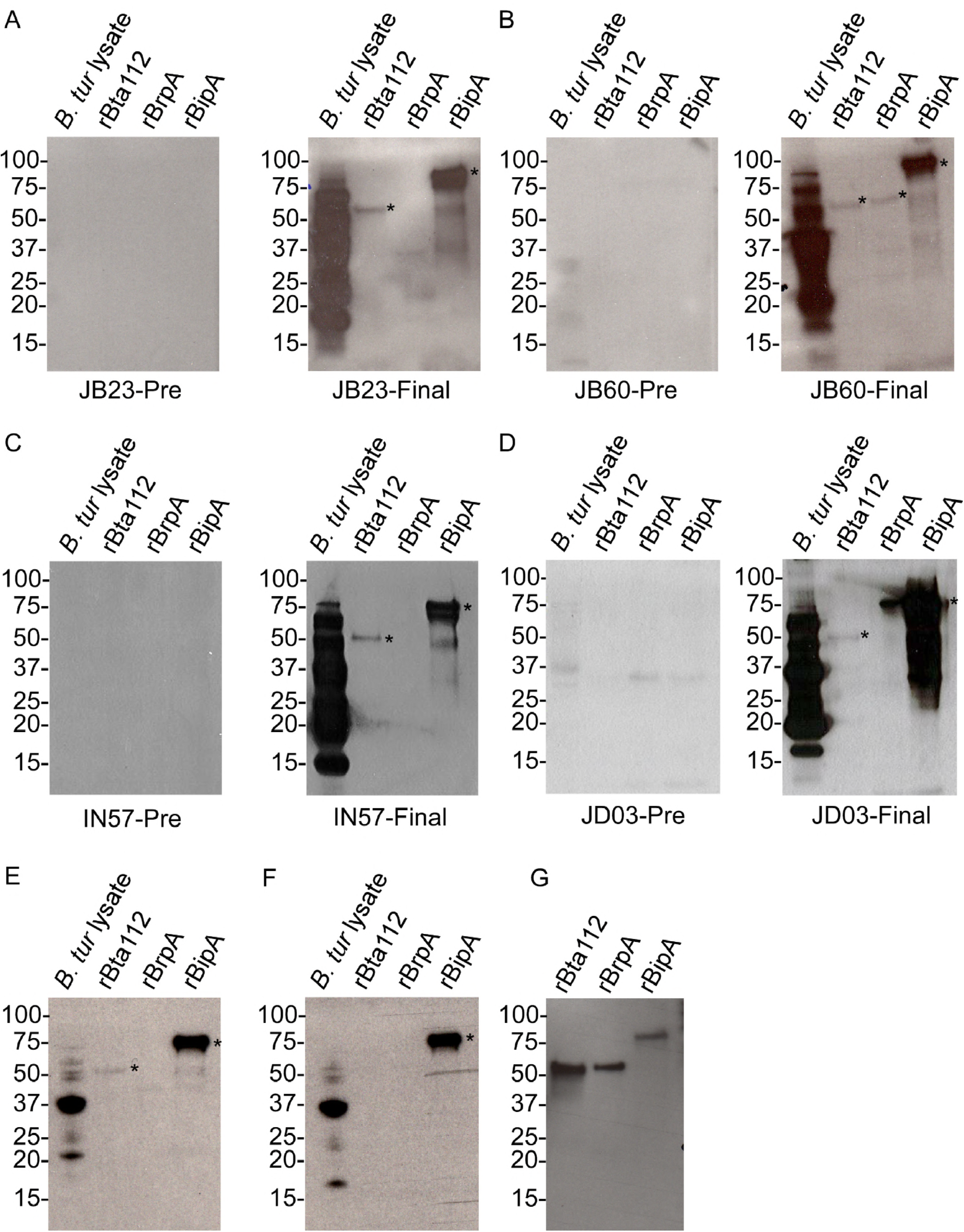
Immunoblot analysis using serum samples from NHPs and mice to rBta112, rBrpA, and rBipA. Immunoblots of JB23 (A), JB60 (B), IN57 (C), and JD03 (D) are shown. Membranes were probed with preinfection serum samples (left blot) and serum samples collected at days 84 (JB23, JB60, and IN57) and 100 (JD03). Immunoblots from two mice (E and F), which represent the remaining 10 animals are shown. Membranes were also probed with an anti-6 histidine monoclonal antibody (G), and indicate the molecular weight of each recombinant protein. An asterisk is next to each recombinant protein that was antigenic. Molecular weights are indicated at the left of each immunoblot.

Since rBipA and rBta112 were immunogenic by immunoblotting in all four NHPs, we further evaluated their serological responses over the duration of the study using enzyme-linked immunosorbent assay (ELISA). Assessment of rBipA and rBta112 indicated the temporal persistence of IgG responses to the recombinant proteins (Figure 6 A-D). JB23, JB60, and IN57 generated IgG responses for at least 84 days after the animals were infected with *B. turicatae* by tick transmission (Figure 6 A-C). These responses were statistically significant compared to the pre-infection serum samples for each animal to a given recombinant protein. JD03 was a control animal, as described in our previous report (46), and evaluating IgG response at three time points (7, 27, and 43 days) after feeding uninfected ticks indicated that tick saliva did not generate cross reactive antibody responses to rBipA and rBta112 (D). After infecting the animal by tick transmission, IgG responses to rBipA and rBta112 were detected 42 days after feeding (D, day 100 of the study). Statistically significant IgG responses were no longer detected to rBta112 from animal JD03 84 days after infection. Collectively, these findings indicated temporal persistence of IgG responses to rBipA, while three of four animals generated prolonged IgG responses to rBta112.

**Figure 6.**
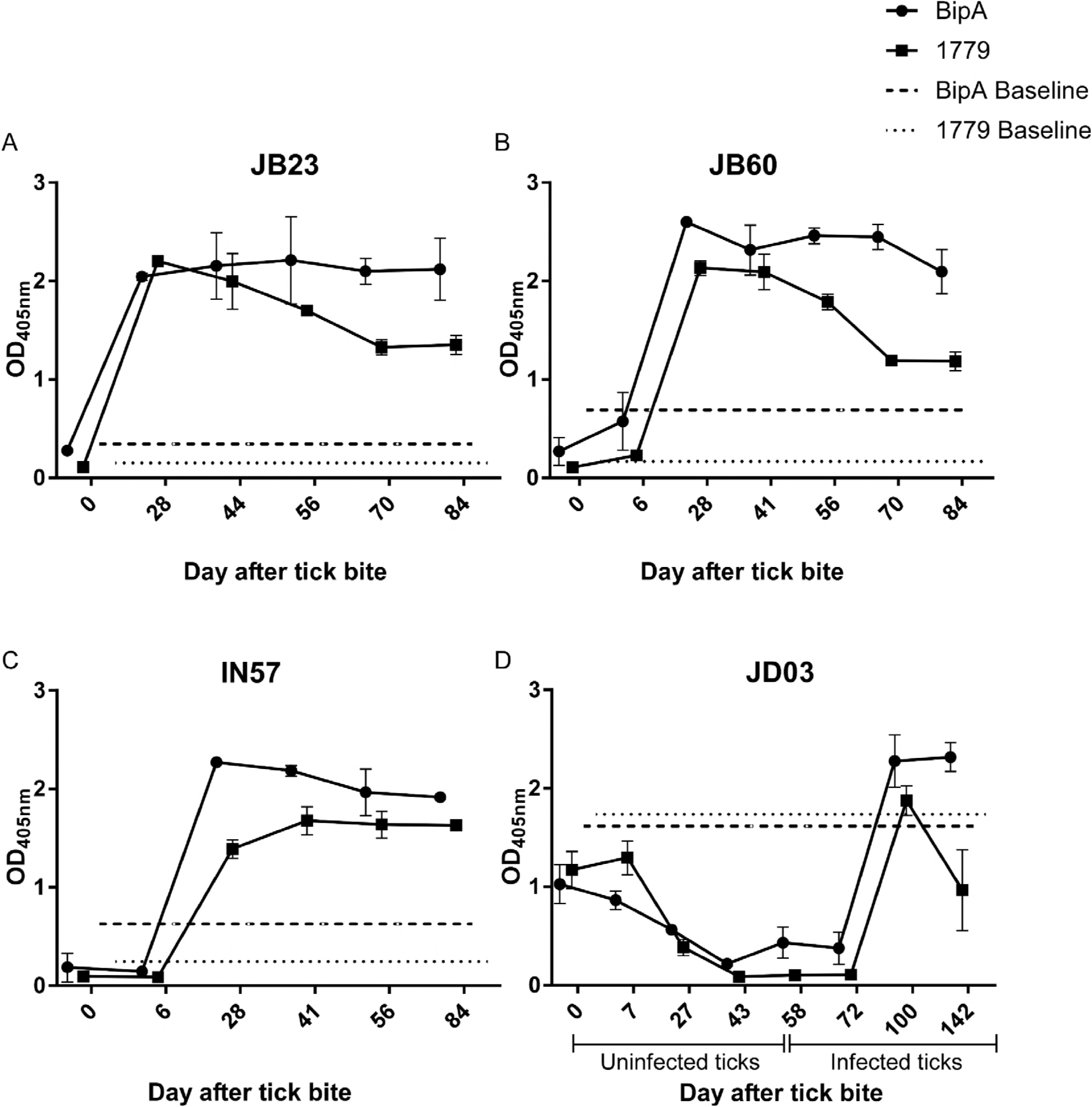
Evaluation of temporal serological responses to rBipA and rBta112 by ELISA. Serum samples were evaluated prior to infection and at days 28, 44, 56, 70, and 84 for animals JB23, JB60, and IN57 (A-C). Serum samples from JD03 were collected prior to and seven, 27, and 43 days after feeding the animal with uninfected (D, uninfected ticks). The animal was then infected with *B. turicatae* by tick bite and serum samples were collected at days 58, 72, 100, and 142 of the study (D, infected ticks). Pre-infection serum samples from each animal were used to establish a statistically significant threshold (p ≤ 0.003) for rBipA (dashed line) and rBta112 (dotted line).

## Discussion

In this study, we identified cytokine profiles associated with pathogenesis and characterized differences in antibody responses of NHPs and mice infected with *B. turicatae*. Evaluating cytokine production from *B*. turicatae-stimulated PBMCs identified mediators involved in disease manifestation. *B. turicatae* induced significant increases in TNFα, IL-1β, sCD40, and IL-23 compared to medium controls. The observed TNFα response has been linked to spirochete lipoprotein-induced Jarisch-Herxheimer reactions [14]. Interestingly, *B. turicatae* also induced statistically significant higher levels of IL-1β compared to cells incubated with *Borrelia burgdorferi*, the Lyme disease pathogen. IL-1β and TNFα are known to play a primary role in triggering miscarriage and pre-term labor in rhesus macaques (47) and in human patients (48, 49). If significant quantities of RF spirochetes cross the placenta, such a response could be induced *in utero*, and this pathogenic mechanism should be further evaluated. We did not detect elevated levels of these key inflammatory mediators directly in serum of infected monkeys; however, the response in PBMCs was detected between 12–24 hours post-stimulation. We therefore suspect that we missed the height of the inflammatory response with evaluation commencing after 14 days of infection. In mice infected with *B. hermsii*, plasma levels of IFN appear to be elevated at the height of spirochetemia, whereas IL-1β is detected after clearance of the infection (50). Directly comparable experiments in mice and primates would be of benefit, but consistent detection of these inflammatory mediators in blood cells exposed to RF spirochetes indicates that they are likely important for pathogenesis.

Our findings suggest unique characteristics in antibody responses generated to *B. turicatae* antigens between mammalian species. BipA is known to be immunogenic in mice (33, 34), but previous work screening serum samples from a small cohort of mice naturally infected with *B. turicatae* by tick bite indicated that BrpA was not antigenic (51). The serological responses from the 10 mice that were evaluated in this current study supported previous findings with BrpA. Interestingly, one NHP produced a detectable response against rBrpA. Furthermore, while only two mice seroconverted to rBta112, the protein was antigenic in all four NHPs. While more animals are needed to definitively determine differences in antibody responses between mice and NHPs, these findings suggested that the immune response between the two mammalian species were dissimilar. Future work should evaluate NHPs as a model for antigen discovery and vaccine development.

B cells drive the immune effort to control infection with relapsing fever spirochetes, and distinct subsets with roles in immunity have been delineated (20, 52, 53). Mature B cells can be divided into follicular (FO) B cells, present in the lymphoid follicles, marginal zone (MZ) B cells, located in the marginal sinus of the spleen, and B1 cells predominantly found in the mouse peritoneum. These are subdivided into B1a and B1b cells. B1 and MZ B cells are known to engage in the T-cell independent antibody response. In contrast to the Lyme disease (LD) spirochete, which induces an expansion of MZ B cells upon infection, RF spirochetes induce a loss of MZ B cells (54). This may reflect the long-term presence of LD spirochetes in the spleen versus the periodic blood-borne expansion of RF spirochetes and/or the differential responses to antigens. The importance of the B1b cell component of the T-independent response to RF spirochetes was demonstrated by transfer of B1b lymphocytes from convalescent mice to Rag 1-/-mice (lacking mature B and T lymphocytes). This subpopulation conferred protection that consisted of a specific IgM response which occurred when mice were challenged 60 days after the reconstitution, indicating that this population alone could confer memory and afford protection (20). Importantly, the identical counterpart of this particular B cell subset has not been identified in humans (55), so it remains to be seen if the same mechanism to control infection occurs in RF patients. Our study shows a precipitous drop in all of the major B cells subsets within the peripheral blood of RF spirochete-infected NHP during the height of bacteremia (2 weeks p.i.). While we did not examine lymph node populations, we surmise that the steep decline in peripheral B cells was met with migration to lymphoid organs. By day 70, no specific B cell subset emerged at an increased frequency over the others. In addition, the specific antibody responses to recombinant proteins were of IgG isotype. Our attempts at screening IgM responses did not produce clear and specific binding to either recombinant proteins or *B.turicatae* lysates. This suggests that B cell responses in primates may rely more on T-dependent IgG subclass responses.

RF Borreliosis is a major burden to maternal and fetal health, especially in resource-poor areas and novel intervention strategies are needed. According to the World Health Organization, every year 45% of all deaths in children under 5 years are among newborn infants in their first 28 days of life or the neonatal period, and 25% of neonatal deaths result from infections (56). The major threat of RF borreliosis caused by both Old and New World species are pregnancy complications occurring during the perinatal period (~20 weeks after gestation to 1–4 weeks after birth) (57–65). The use of rhesus macaques as an animal model resulted in the identification of a novel antigens (BrpA and Bta112) and confirmed the immunogenicity of BipA. Future studies will determine if these antigens offer potential targets for human vaccination or in diagnosis of RF. We will also expand on the developed NHP model and focus on understanding immunopathology of infected pregnant macaques to identify cytokines in amniotic fluid, which may reveal a mechanism of perinatal effects associated with RF infection.

## Materials and Methods

### Ethics statement

Practices in the housing and care of NHP and mice conformed to the regulations and standards of the Public Health Service Policy on Humane Care and Use of Laboratory Animals, and the Guide for the Care and Use of Laboratory Animals. The Tulane National Primate Research Center (TNPRC) and Baylor College of Medicine (BCM) are fully accredited by the Association for the Assessment and Accreditation of Laboratory Animal Care-International. The Institutional Animal Care and Use Committees at the TNPRC and BCM approved all animal-related protocols, including the infection and sample collection from NHPs and mice. All animal procedures were overseen by veterinarians and their staff.

### *B. turicatae* strains used and animal infections by tick bite

*B. turicatae* strains used in this study were 91E135, Florida canine *Borrelia* (FCB), 99PE-1807, Texas canine *Borrelia* (TCB) 1, and TCB2 (66). Tick transmission studies to were previously reported using a colony of *O. turicata* that originated from Kansas (31). Briefly, four male Indian rhesus macaques (JB23, JB60, IN57, and JD03) 2.02–2.85 years of age were used. Animals were sedated with 5–8 mg/kg Telazol by intramuscular injection and ten third stage nymphal ticks infected with *B. turicatae* were fed on each NHP (67). JD03 was initially fed upon by 10 uninfected ticks and monitored for 42 days as a control for tick-specific responses. This animal was subsequently fed upon by infected ticks.

Murine infection by tick bite was performed as previously described (34). Eight to 10 infected third stage nymphal *O. turicata* were fed to repletion on 10 Institute of Cancer Research (ICR) mice, a Swiss derivative maintained at BCM. Infection was assessed by collecting a drop of blood from the animals and evaluating the specimen by dark field microscopy for the presence of circulating spirochetes. Thirty days after infection by tick bite, the animals were exsanguinated and serum samples were obtained.

### Collection and processing of NHP blood

To evaluate immune responses from NHPs, both whole blood and clotted blood for serum were collected. Animals were anesthetized (Ketamine, 0.1 ml/kg, IM) and blood was collected by venipuncture of the femoral vein into either clot tubes or EDTA tubes (whole blood). Blood for serum samples was collected at day 0 (prior to tick feeding), day 28, day 56, day 70 and day 85, as previously described (31). Whole blood for flow cytometry was collected at day 0, day 14 and day 70. Tubes containing clotted blood were centrifuged at 3,000 rpm for 10 minutes to obtain serum samples. Peripheral blood mononuclear cells (PBMCs) were isolated from whole blood using Lymphocyte Separation Medium (MP Biomedicals) (68). The lymphocyte layer was washed once with sterile PBS, then resuspended in PBS/2% FBS and counted. Cells were again pelleted and resuspended in Freeze Medium (Invitrogen) at ≤1 × 10^7^/ml, then cryopreserved in liquid nitrogen until staining.

### Flow cytometry Assay

Cryopreserved PBMCs were thawed, washed in RPMI-1640 media, counted with trypan blue exclusion staining, and adjusted to a concentration of 1 × 10^7^ cells/ml in RPMI-1640 media with 10% FBS. One hundred μl of cells were used for staining with different concentrations of monoclonal antibodies and incubated for 25 min at room temperature, protected from light, as reported earlier (68–71). The cells were further washed two times with 3ml of flow wash buffer (PBS with 0.1% BSA and 7mM sodium azide) and centrifuged at 1350rpm for 7 min. Following aspiration of supernatants from cell pellets, the cell pellets were resuspended in 350 μl of 1% paraformaldehyde buffer (in PBS). For antibodies conjugated with tandem dyes, the cell pellets were dissolved in FACS fixation and stabilization buffer (Becton Dickinson). For T cell phenotyping, CD3-FITC (SP34–2, BD Biosciences), CD8-PerCP (SK1, BD Biosciences), and CD4-APC (L200, BD Biosciences) were used. For B cell phenotyping, anti-CD5-PE-Cy5.5 (CD5–5D7, Invitrogen), anti-CD20-ECD (B9E9, Beckman Coulter), anti-CD21-APC (B-Ly4, BD Biosciences), anti-CD86-PECy5 (FUN-1, BD Biosciences) anti-CD138-FITC (MI15, BD Biosciences), anti-CD27-FITC (M-T271 BD Biosciences), anti-IgM (G20-127, BD Biosciences) and anti-IgD (purified polyclonal, Southern Biotech) antibodies were used. Anti-CD38 antibody (clone OKT10) was obtained from NIH NHP Reagent Resource. Data were acquired within 24 hours of staining using either BD Fortessa instrument (BD Immunocytometry System) or BD Facsverse (BD Biosciences) and FACSDiva software (BD Immunocytometry System). For each sample, 50,000 events were collected by gating either on CD3+ T cells or CD20+ B cells. For B-cell phenotypic analysis, cells were first gated on singlets, followed by lymphocytes, and CD20+ B-cells and CD20-cells. CD20+ B-cells were further gated for CD5/CD21/CD86 and CD138 expression. In cases where enough PBMCs were available, flow cytometry was repeated to give duplicate samples. The gating strategy, along with representative results from a single animal, is shown in supplementary Figure S1.

### Cytokine/chemokine array

A portion of PBMCs derived from whole blood were also used for in vitro stimulation with *B. turicatae*. Cells isolated from blood collected from each animal (which included JD03 following control/uninfected tick feeding) on day 14 post-tick feeding were resuspended in RPMI 1640/10% FBS at 1 × 10^6^/ml and 0.5 ml was added to each well of a 24-well plate. Late log-phase *B. turicatae* was diluted to 1 × 10^7^ spirochetes per ml and 0.5 ml was added to appropriate wells for a 10:1 ratio of spirochetes to cells. Controls included untreated cells, cells incubated with *B. burgdorferi*, and cells incubated with BSK medium (Sigma). To determine the impact of soluble factors produced by the spirochetes, cells were incubated with 0.22 μm-filtered BSK medium derived from *B. turicatae* and *B. burgdorferi* cultures. Cultures were placed in a 37°C, 5% CO_2_ incubator. Supernatants were collected at 12 and 24 hours and stored at −20°C. Serum samples from days 14, 28 and 41 were also tested by the cytokine/chemokine array. Undiluted samples were analyzed using the MILLIPLEX MAP Non-Human Primate Cytokine Magnetic Bead Panel - Premixed 23 Plex (Millipore) according to the manufacturer’s instructions. The bead assay was performed by the Pathogen Detection and Quantification Core at the TNPRC and analyzed on a Bioplex 2000 Suspension Array System (BioRad). Each analyte concentration was calculated by logistic-5PL regression of the standard curve. To determine the statistical significance between the means for two experimental groups, an unpaired, two-tailed Student’s t-test was performed using GraphPad Software QuickCalcs. Those differences with p≤ 0.05 are reported as significant.

### Computational analysis of Bta112

Initially, *bta112* was identified as a gene up-regulated by *B. turicatae* in the tick and at 22°C (tick-like growth conditions) compared to spirochetes isolated from infected murine blood and spirochete grown at 35°C (mammalian-like growth conditions) (43). The protein was evaluated using the Basic Local Alignment Search Tool (BLAST) from NCBI, LipoP1.0, and ScanProsite. The gene sequence of *bta112* was evaluated in *B. turicatae* 91E135, FCB, 99PE-1807, TCB 1, and TCB 2 through ongoing genome sequencing efforts of these isolates.

### Recombinant proteins and rabbit serum generation to recombinant Bta112 (rBta112)

Recombinant BipA (rBipA) and BrpA (rBrpA) were produced as six histidine linked proteins as previously described (34, 51). Recombinant Bta112 was also produced as a six histidine linked recombinant protein using the pEXP1-DEST expression vector (ThermoFisher Scientific, Waltham, MA). The *bta112* gene was amplified by PCR from *B. turicatae* gDNA with Accuprime Pfx (Thermo Fisher Scientific) without its predicted signal sequence (1–69 bp, signal P 3.0 -http://www.cbs.dtu.dk/services/SignalP-3.0/). The gene’s signal sequence was omitted from amplification using primers SP/1779/70–1461 (5'-CAAACAAGTTTGTACAAAAATTTCAAAAGTCCAAAAGACGCTG-3') and ASP/1779/70–1461 (5'-CGTATGGGTAAAGC TTATTACTACTTGCGGTACTATCTGCTG-3'). The amplicon was cloned by In-fusion (BD Clontech) into pEXP1-HA-DEST, digested with BsrGI and HindIII to create pEXP1-HA::bta112, and Top10 *Escherichia coli* were transformed. Plasmid DNA was isolated and submitted for sequencing to ensure that errors were not introduced by PCR. Vector NTI 11.0 (ThermoFisher Scientific) was used to assess *bta112* sequence. rBta112 was produced by transforming *E. coli* BL21 Star (DE3) cells (ThermoFisher Scientific) with pEXP1-HA::bta112 and expression was induced with 0.5 mM IPTG. rBta112 was purified by nickel chelate chromatography.

Rabbit anti-rBta112 was produced by Cocalico Biologicals, INC. Pre-immunization serum samples were collected from two rabbits and the animals were immunized intraperitoneally with 50 μg of rBta112 using complete Freund’s adjuvant. The animals were immunized three subsequent times in two-week intervals using Freund’s incomplete adjuvant. Serum samples were collected and evaluated for specificity to rBta112 and the native protein by immunoblotting.

### Surface localization assays, immunoblotting, and densitometry analysis

To determine the surface localization of Bta112, proteinase K assays and immunoblotting were performed as previously described (51, 72). Moreover, for all immunoblotting assays *B. turicatae* was grown at 35°C. For proteinase K assays, spirochetes were grown to a density > 5 × 10^7^ cells per ml, pelleted at 1,000 × *g* for 10 minutes at room temperature, washed in PBS + MgCl_2_, pelleted again, and resuspended in PBS + MgCl_2_. Spirochetes were incubated with increasing concentrations (5, 50, and 200 μg per ml) of proteinase K (Promega, Madison, WI) for 15 minutes at room temperature. PBS + MgCl_2_ was used as a vehicle control. Proteinase K was inactivated by boiling samples at 100°C for 10 minutes. SDS-PAGE and immunoblotting were performed as previously described using the Any kD Mini-PROTEAN TGX Stain-free precast gels (BioRad, Hercules, CA) (51). One μg of recombinant protein or 1 × 10^7^ spirochetes were electrophoresed per lane, and the Trans-Blot Cell (BioRad) was used to transfer proteins onto polyvinylidene fluoride membranes. Rabbit, murine, chicken, and NHP serum samples were used to probe immunoblots at a concentration of 1:200, and antibody binding was detected with the appropriate secondary antibody and the ECL Western blotting reagent (VWR, Atlanta, GA). ImageLab (6.0.1) was used to quantify the relative density of FlaB of spirochetes incubated with 5, 50 and 200 μg per ml proteinase K to spirochetes that did not undergo proteinase K treatment.

### ELISA

Immulon 2HB flat bottom microtiter polystyrene plates (Thermo Fisher, Waltham, MA) were coated with 1 μg/ml of rBipA or r1779 using 1x coating solution (KPL, Gaithersburg, MD). The plates were washed three times with wash buffer (1x PBS and 0.05% Tween20) and blocked with diluent (1x PBS, 5% Horse Serum, 0.05% Tween20, 0.001% Dextran Sulfate) overnight at 4°C. Plates were washed again and probed with the NHP serum samples at a 1:100 dilution in diluent and incubated for one hour at room temperature. Plates were washed again and incubated for one hour at room temperature with peroxidase labeled goat anti-monkey IgG (KPL, Gaithersburg, MD) at a 1:4000 dilution. Plates were washed again and incubated with ABTS Peroxidase Substrate (KPL, Gaithersburg, MD) for 30 minutes and read at 405nm on an Epoch Microplate Spectrophotometer (Biotek, Winooski, VT). Samples were considered statistically significant if their mean optical density was more than three times the SD above the mean of the pre-tick challenge sera (p ≤ 0.003).

**Table 1.**
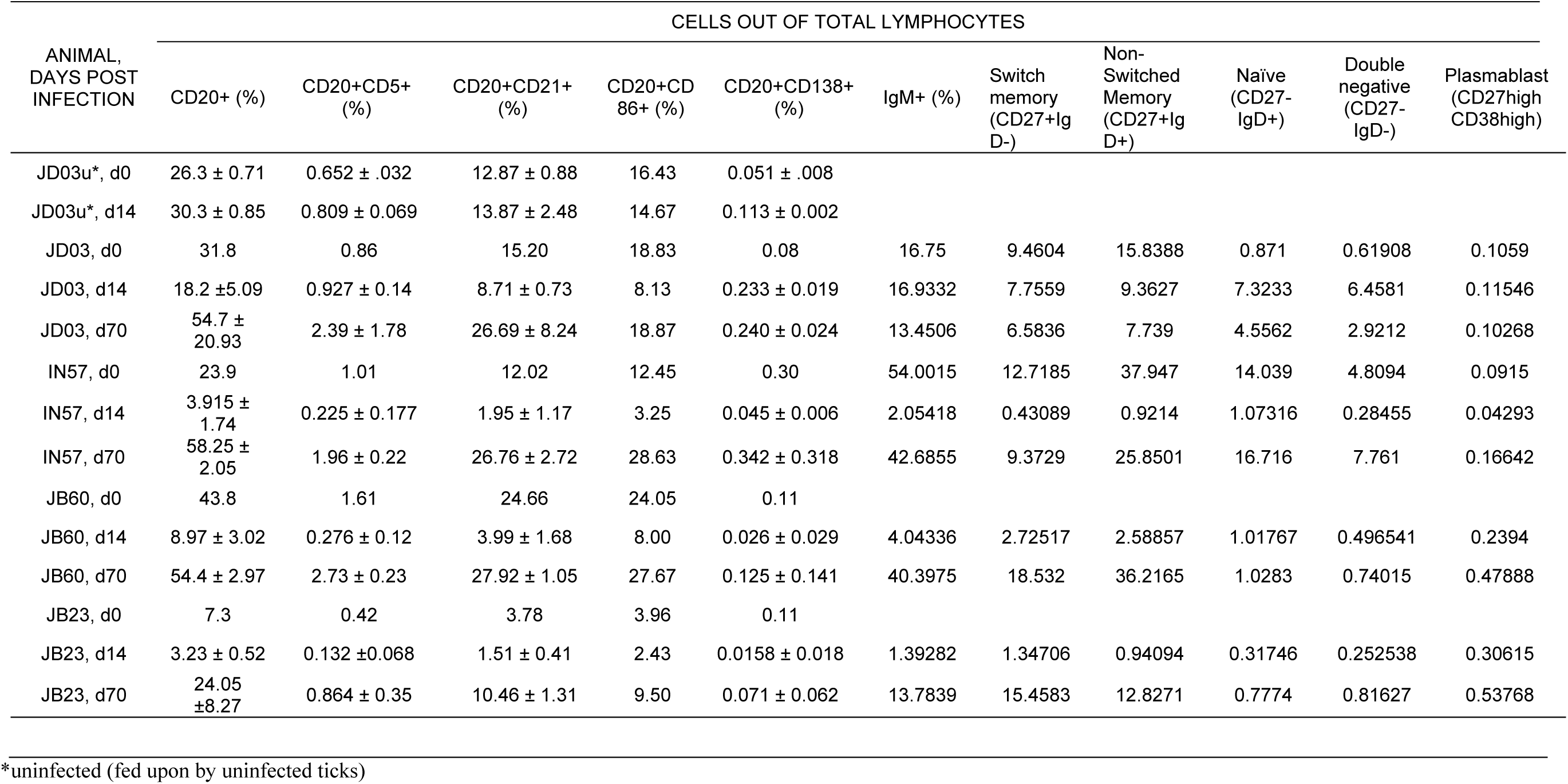
Changes in Peripheral B cell Subsets Following Infection with *B. turicatae*.

| ANIMAL,<br>DAYS POST<br>INFECTION | CELLS OUT OF TOTAL LYMPHOCYTES |  |  |  |  |  |  |  |  |  |  |
| --- | --- | --- | --- | --- | --- | --- | --- | --- | --- | --- | --- |
|  | CD20+ (%) | CD20+CD5+<br>(%) | CD20+CD21+<br>(%) | CD20+CD<br>86+ (%) | CD20+CD138+<br>(%) | IgM+ (%) | Switch<br>memory<br>(CD27+Ig<br>D-) | Non-<br>Switched<br>Memory<br>(CD27+Ig<br>D+) | Naïve<br>(CD27-<br>IgD+) | Double<br>negative<br>(CD27-<br>IgD-) | Plasmablast<br>(CD27high<br>CD38high) |
| JD03u*, d0 | 26.3 ± 0.71 | 0.652 ± .032 | 12.87 ± 0.88 | 16.43 | 0.051 ± .008 |  |  |  |  |  |  |
| JD03u*, d14 | 30.3 ± 0.85 | 0.809 ± 0.069 | 13.87 ± 2.48 | 14.67 | 0.113 ± 0.002 |  |  |  |  |  |  |
| JD03, d0 | 31.8 | 0.86 | 15.20 | 18.83 | 0.08 | 16.75 | 9.4604 | 15.8388 | 0.871 | 0.61908 | 0.1059 |
| JD03, d14 | 18.2 ± 5.09 | 0.927 ± 0.14 | 8.71 ± 0.73 | 8.13 | 0.233 ± 0.019 | 16.9332 | 7.7559 | 9.3627 | 7.3233 | 6.4581 | 0.11546 |
| JD03, d70 | 54.7 ±<br>20.93 | 2.39 ± 1.78 | 26.69 ± 8.24 | 18.87 | 0.240 ± 0.024 | 13.4506 | 6.5836 | 7.739 | 4.5562 | 2.9212 | 0.10268 |
| IN57, d0 | 23.9 | 1.01 | 12.02 | 12.45 | 0.30 | 54.0015 | 12.7185 | 37.947 | 14.039 | 4.8094 | 0.0915 |
| IN57, d14 | 3.915 ±<br>1.74 | 0.225 ± 0.177 | 1.95 ± 1.17 | 3.25 | 0.045 ± 0.006 | 2.05418 | 0.43089 | 0.9214 | 1.07316 | 0.28455 | 0.04293 |
| IN57, d70 | 58.25 ±<br>2.05 | 1.96 ± 0.22 | 26.76 ± 2.72 | 28.63 | 0.342 ± 0.318 | 42.6855 | 9.3729 | 25.8501 | 16.716 | 7.761 | 0.16642 |
| JB60, d0 | 43.8 | 1.61 | 24.66 | 24.05 | 0.11 |  |  |  |  |  |  |
| JB60, d14 | 8.97 ± 3.02 | 0.276 ± 0.12 | 3.99 ± 1.68 | 8.00 | 0.026 ± 0.029 | 4.04336 | 2.72517 | 2.58857 | 1.01767 | 0.496541 | 0.2394 |
| JB60, d70 | 54.4 ± 2.97 | 2.73 ± 0.23 | 27.92 ± 1.05 | 27.67 | 0.125 ± 0.141 | 40.3975 | 18.532 | 36.2165 | 1.0283 | 0.74015 | 0.47888 |
| JB23, d0 | 7.3 | 0.42 | 3.78 | 3.96 | 0.11 |  |  |  |  |  |  |
| JB23, d14 | 3.23 ± 0.52 | 0.132 ± 0.068 | 1.51 ± 0.41 | 2.43 | 0.0158 ± 0.018 | 1.39282 | 1.34706 | 0.94094 | 0.31746 | 0.252538 | 0.30615 |
| JB23, d70 | 24.05<br>± 8.27 | 0.864 ± 0.35 | 10.46 ± 1.31 | 9.50 | 0.071 ± 0.062 | 13.7839 | 15.4583 | 12.8271 | 0.7774 | 0.81627 | 0.53768 |

\*uninfected (fed upon by uninfected ticks)

## Acknowledgements

Funding for this work was provided by a TNPRC Pilot Grant (MEE and JL), NIH grants AI091652 and AI103724 (JL), and the TNPRC base grant (NIH) 5 P51 OD 011104-56. The authors would like to thank Dr. Britton Grasperge (Louisiana State University), and Amanda C. Tardo (TNPRC) for their assistance with experiments.

## References

1. Dworkin MS, Schwan TG, Anderson DE, Jr., Borchardt SM. 2008. Tick-borne relapsing fever. Infect Dis Clin North Am 22:449–468, viii.

2. Fukunaga M, Koreki Y. 1995. The flagellin gene of *Borrelia miyamotoi* sp. nov. and its phylogenetic relationship among *Borrelia* species. FEMS Microbiol Lett 134: 255–258.

3. Geller J, Nazarova L, Katargina O, Jarvekulg L, Fomenko N, Golovljova I. 2012. Detection and genetic characterization of relapsing fever spirochete Borrelia miyamotoi in Estonian ticks. PLoS One 7:e51914.

4. Breuner NE, Dolan MC, Replogle AJ, Sexton C, Hojgaard A, Boegler KA, Clark RJ, Eisen L. 2017. Transmission of *Borrelia miyamotoi* sensu lato relapsing fever group spirochetes in relation to duration of attachment by *Ixodes scapularis* nymphs. Ticks Tick Borne Dis 8: 677–681.

5. Melkert PW. 1987. Fatal-Jarisch Herxheimer reaction in a case of relapsing fever misdiagnosed as lobar pneumonia. Tropical & Geographical Medicine 39: 92–93.

6. Rustenhoven-Spaan I, Melkert P, Nelissen E, Roosmalen Jv, Stekelenburg J. 2013. Maternal mortality in a rural Tanzanian hospital: fatal Jarisch-Herxheimer reaction in a case of relapsing fever in pregnancy. Tropical Doctor 43: 138–141.

7. Y Negussie DGR, L E DeForge, S L Kunkel, A Eynon, G E Griffin. 1992. Detection of plasma tumor necrosis factor, interleukins 6, and 8 during the Jarisch-Herxheimer Reaction of relapsing fever. The Journal of Experimental Medicine 175: 1207–1212.

8. Melkert PW. 1991. Mortality in high risk patients with tick-borne relapsing fever analysed by the Borrelia-index. East Afr Med J 68: 875–879.

9. Melkert PW, Stel HV. 1991. Neonatal Borrelia infections (relapsing fever): report of 5 cases and review of the literature. East African Medical Journal 68: 999–1005.

10. Barclay AJ, Coulter JB. 1990. Tick-borne relapsing fever in central Tanzania. Trans R Soc Trop Med Hyg 84: 852–856.

11. McConnell J. 2003. Tick-borne relapsing fever under-reported. The Lancet Infectious Diseases 3: 604.

12. Larsson C, Andersson M, Bergstrom S. 2009. Current issues in relapsing fever. Current Opinion in Infectious Diseases 22: 443–449.

13. Lundqvist J, Larsson C, Nelson M, Andersson M, Bergström S, Persson C. 2010. Concomitant Infection Decreases the Malaria Burden but Escalates Relapsing Fever Borreliosis. Infection and Immunity 78: 1924–1930.

14. Nordstrand A, Bunikis I, Larsson C, Tsogbe K, Schwan TG, Nilsson M, Bergstrom S. 2007. Tickborne relapsing fever diagnosis obscured by malaria, Togo. Emerging Infectious Diseases 13: 117–123.

15. Yokota M, Morshed MG, Nakazawa T, Konishi H. 1997. Protective activity of Borrelia duttonii-specific immunoglobulin subclasses in mice. Journal of Medical Microbiology 46: 675–680.

16. Alugupalli KR, Gerstein RM, Chen J, Szomolanyi-Tsuda E, Woodland RT, Leong JM. 2003. The resolution of relapsing fever borreliosis requires IgM and is concurrent with expansion of B1b lymphocytes. Journal of Immunology 170: 3819–3827.

17. Newman K, Johnson RC. 1984. T-cell-independent elimination of Borrelia turicatae. Infection and Immunity 45: 572–576.

18. Barbour AG, Bundoc V. 2001. In vitro and in vivo neutralization of the relapsing fever agent Borrelia hermsii with serotype-specific immunoglobulin M antibodies. Infection & Immunity 69: 1009–1015.

19. Connolly SE, Benach JL. 2001. Cutting edge: the spirochetemia of murine relapsing fever is cleared by complement-independent bactericidal antibodies. Journal of Immunology 167: 3029–3032.

20. Alugupalli KR, Leong JM, Woodland RT, Muramatsu M, Honjo T, Gerstein RM. 2004. B1b lymphocytes confer T cell-independent long-lasting immunity. Immunity 21: 379–390.

21. Malkiel S, Kuhlow CJ, Mena P, Benach JL. 2009. The loss and gain of marginal zone and peritoneal B cells is different in response to relapsing fever and Lyme disease Borrelia. Journal of Immunology 182: 498–506.

22. Boyle WK, Wilder HK, Lawrence AM, Lopez JE. 2014. Transmission dynamics of *Borrelia turicatae* from the arthropod vector. PLoS Negl Trop Dis 8:e2767.

23. Schwan TG, Hinnebusch BJ. 1998. Bloodstream-versus tick-associated variants of a relapsing fever bacterium. Science 280: 1938–1940.

24. Krajacich BJ, Lopez JE, Raffel SJ, Schwan TG. 2015. Vaccination with the variable tick protein of the relapsing fever spirochete *Borrelia hermsii* protects mice from infection by tick-bite. Parasit Vectors 8: 546.

25. Bennett IL, Jr., Cluff LE. 1957. Bacterial pyrogens. Pharmacol Rev 9: 427–479.

26. Colvin JW, Mills CA. 1939. Ease of Body Heat Loss and Resistance to Infection. Science 90: 275–276.

27. Halberg F, Spink WW. 1956. The influence of Brucella somatic antigen (endotoxin) upon the temperature rhythm of intact mice. Lab Invest 5: 283–294.

28. Wright DJ, Woodrow DF. 1980. Terminal changes in mice experimentally infected with Borrelia duttoni. J Pathol 130: 83–90.

29. Klein MS, Conn CA, Kluger MJ. 1992. Behavioral thermoregulation in mice inoculated with influenza virus. Physiol Behav 52: 1133–1139.

30. Francis E. 1938. Longevity of the tick Ornithodoros turicata and of *Spirochaeta recurrentis* within this tick. Publ Hlth Rep:2220–2241.

31. Lopez JE, Vinet-Oliphant H, Wilder HK, Brooks CP, Grasperge BJ, Morgan TW, Stuckey KJ, Embers ME. 2014. Real-Time Monitoring of Disease Progression in Rhesus Macaques Infected with *Borrelia turicatae* by Tick Bite. J Infect Dis doi:10.1093/infdis/jiu306.

32. Seok J, Warren HS, Cuenca AG, Mindrinos MN, Baker HV, Xu W, Richards DR, McDonald-Smith GP, Gao H, Hennessy L, Finnerty CC, López CM, Honari S, Moore EE, Minei JP, Cuschieri J, Bankey PE, Johnson JL, Sperry J, Nathens AB, Billiar TR, West MA, Jeschke MG, Klein MB, Gamelli RL, Gibran NS, Brownstein BH, Miller-Graziano C, Calvano SE, Mason PH, Cobb JP, Rahme LG, Lowry SF, Maier RV, Moldawer LL, Herndon DN, Davis RW, Xiao W, Tompkins RG, Inflammation t, Host Response to Injury LSCRP. 2013. Genomic responses in mouse models poorly mimic human inflammatory diseases. Proceedings of the National Academy of Sciences 110: 3507–3512.

33. Lopez JE, Schrumpf ME, Nagarajan V, Raffel SJ, McCoy BN, Schwan TG. 2010. A novel surface antigen of relapsing fever spirochetes can discriminate between relapsing fever and Lyme borreliosis. Clin Vaccine Immunol 17: 564–571.

34. Lopez JE, Wilder HK, Boyle W, Drumheller LB, Thornton JA, Willeford B, Morgan TW, Varela-Stokes A. 2013. Sequence analysis and serological responses against *Borrelia turicatae* BipA, a putative species-specific antigen. PLoS Negl Trop Dis 7:e2454.

35. Yammani RD, Haas KM. 2013. Primate B-1 Cells Generate Antigen-Specific B Cell Responses to T Cell-Independent Type 2 Antigens. The Journal of Immunology 190: 3100–3108.

36. Tung JW, Mrazek MD, Yang Y, Herzenberg LA, Herzenberg LA. 2006. Phenotypically distinct B cell development pathways map to the three B cell lineages in the mouse. Proceedings of the National Academy of Sciences 103: 6293–6298.

37. Perez-Andres M, Paiva B, Nieto WG, Caraux A, Schmitz A, Almeida J, Vogt RF, Marti GE, Rawstron AC, Van Zelm MC, Van Dongen JJM, Johnsen HE, Klein B, Orfao A, and the Primary Health Care Group of Salamanca for the Study of MBL. 2010. Human peripheral blood B-cell compartments: A crossroad in B-cell traffic. Cytometry Part B: Clinical Cytometry 78B:S47–S60.

38. Roozendaal R, Carroll MC. 2007. Complement receptors CD21 and CD35 in humoral immunity. Immunological Reviews 219: 157–166.

39. Das A, Xu H, Wang X, Yau CL, Veazey RS, Pahar B. 2011. Double-Positive CD21+CD27+ B Cells Are Highly Proliferating Memory Cells and Their Distribution Differs in Mucosal and Peripheral Tissues. PLoS ONE [Electronic Resource] 6:e16524.

40. Neumann B, Sopper S, Stahl-Hennig C. 2015. OMIP-026: Phenotypic analysis of B and plasma cells in rhesus macaques. Cytometry Part A 87: 800–802.

41. O'Neill SK, Cao Y, Hamel KM, Doodes PD, Hutas G, Finnegan A. 2007. Expression of CD80/86 on B Cells Is Essential for Autoreactive T Cell Activation and the Development of Arthritis. The Journal of Immunology 179: 5109.

42. Bar-Or A, Oliveira EML, Anderson DE, Krieger JI, Duddy M, O'Connor KC, Hafler DA. 2001. Immunological Memory: Contribution of Memory B Cells Expressing Costimulatory Molecules in the Resting State. The Journal of Immunology 167: 5669.

43. Wilder HK, Raffel SJ, Barbour AG, Porcella SF, Sturdevant DE, Vaisvil B, Kapatral V, Schmitt DP, Schwan TG, Lopez JE. 2016. Transcriptional Profiling the 150 kb Linear Megaplasmid of *Borrelia turicatae* Suggests a Role in Vector Colonization and Initiating Mammalian Infection. PLoS One 11:e0147707.

44. Bekeredjian-Ding I, Jego G. 2009. Toll-like receptors - sentries in the B-cell response. Immunology 128: 311–323.

45. Liang FT, Philipp MT. 1999. Analysis of antibody response to invariable regions of VlsE, the variable surface antigen of Borrelia burgdorferi. Infection & Immunity 67: 6702–6706.

46. Lopez JE, Vinet-Oliphant H, Wilder HK, Brooks CP, Grasperge BJ, Morgan TW, Stuckey KJ, Embers ME. 2014. Real-time monitoring of disease progression in rhesus macaques infected with Borrelia turicatae by tick bite. Journal of Infectious Diseases 210: 1639–1648.

47. Sadowsky DW, Adams KM, Gravett MG, Witkin SS, Novy MJ. 2006. Preterm labor is induced by intraamniotic infusions of interleukin-1β and tumor necrosis factor-α but not by interleukin-6 or interleukin-8 in a nonhuman primate model. American Journal of Obstetrics and Gynecology 195: 1578–1589.

48. Vitoratos N, Papadias C, Economou E, Makrakis E, Panoulis C, Creatsas G. 2006. Elevated circulating IL-1beta and TNF-alpha, and unaltered IL-6 in first-trimester pregnancies complicated by threatened abortion with an adverse outcome. Mediators of Inflammation 2006: 30485.

49. Shaarawy M, Nagui AR. 1997. Enhanced expression of cytokines may play a fundamental role in the mechanisms of immunologically mediated recurrent spontaneous abortion. Acta Obstetricia et Gynecologica Scandinavica 76: 205–211.

50. Crowder C, Ghalyanchi Langeroudi A, Shojaee Estabragh A, Lewis E, Marcsisin R, Barbour A. 2016. Pathogen and Host Response Dynamics in a Mouse Model of Borrelia hermsii Relapsing Fever. Veterinary Sciences 3: 19.

51. Lopez JE, Wilder HK, Hargrove R, Brooks CP, Peterson KE, Beare PA, Sturdevant DE, Nagarajan V, Raffel SJ, Schwan TG. 2013. Development of genetic system to inactivate a *Borrelia turicatae* surface protein selectively produced within the salivary glands of the arthropod vector. PLoS Negl Trop Dis 7:e2514.

52. Vaughan AT, Roghanian A, Cragg MS. 2011. B cells—Masters of the immunoverse. The International Journal of Biochemistry & Cell Biology 43: 280–285.

53. Connolly SE, J.L. Benach. 2001. Cutting edge: the spirochetemia of murine relapsing fever is cleared by complement-independent bactericidal antibodies. J Immunol 167: 3029–3032.

54. Malkiel S, Kuhlow CJ, Mena P, Benach JL. 2009. The Loss and Gain of Marginal Zone and Peritoneal B Cells Is Different in Response to Relapsing Fever and Lyme Disease Borrelia. The Journal of Immunology 182: 498–506.

55. Quách TD, Rodríguez-Zhurbenko N, Hopkins TJ, Guo X, Vázquez AMH, Li W, Rothstein TL. 2016. Distinctions Among Circulating Antibody Secreting Cell Populations, Including B-1 Cells, in Human Adult Peripheral Blood(1). Journal of immunology (Baltimore, Md: 1950) 196: 1060–1069.

56. WHO. http://www.who.int/mediacentre/factsheets/fs333/en/. Accessed

57. Dworkin MS, Schwan TG, Anderson DE. 2002. Tick-borne relapsing fever in North America. Med Clin North Amer 86: 417–433.

58. Walker RL, Read DH, Hayes DC, Nordhausen RW. 2002. Equine abortion associated with the *Borrelia parkeri-B. turicatae* tick-borne relapsing fever spirochete group. J Clin Microbiol 40: 1558–1562.

59. Fuchs PC, Oyama AA. 1969. Neonatal relapsing fever due to transplacental transmission of borrelia. JAMA 208: 690–692.

60. Larsson C, Andersson M, Guo BP, Nordstrand A, Hagerstrand I, Carlsson S, Bergstrom S. 2006. Complications of pregnancy and transplacental transmission of relapsing-fever borreliosis. J Infect Dis 194: 1367–1374.

61. Melkert PW. 1987. Prognostic value of the Borrelia-index in relapsing fever. East Afr Med J 64: 284–286.

62. Melkert PW. 1987. Fatal-Jarisch Herxheimer reaction in a case of relapsing fever misdiagnosed as lobar pneumonia. Trop Geogr Med 39: 92–93.

63. Melkert PW, Stel HV. 1991. Neonatal Borrelia infections (relapsing fever): report of 5 cases and review of the literature. East Afr Med J 68: 999–1005.

64. Melkert PW. 1988. Relapsing fever in pregnancy: analysis of high-risk factors. Br J Obstet Gynaecol 95: 1070–1072.

65. Jongen VH, van Roosmalen J, Tiems J, Van Holten J, Wetsteyn JC. 1997. Tick-borne relapsing fever and pregnancy outcome in rural Tanzania. Acta Obstet Gynecol Scand 76: 834–838.

66. Schwan TG, Raffel SJ, Schrumpf ME, Policastro PF, Rawlings JA, Lane RS, Breitschwerdt EB, Porcella SF. 2005. Phylogenetic analysis of the spirochetes *Borrelia parkeri* and *Borrelia turicatae* and the potential for tick-borne relapsing fever in Florida. J Clin Microbiol 43: 3851–3859.

67. Lopez JE, Vinet-Oliphant H, Wilder HK, Brooks CP, Grasperge BJ, Morgan TW, Stuckey KJ, Embers ME. 2014. Real-Time Monitoring of Disease Progression in Rhesus Macaques Infected With Borrelia turicatae by Tick Bite. Journal of Infectious Diseases 210: 1639–1648.

68. Pahar B, Lackner AA, Veazey RS. 2006. Intestinal double-positive CD4+CD8+ T cells are highly activated memory cells with an increased capacity to produce cytokines. European Journal of Immunology 36: 583–592.

69. Pahar B, Gray WL, Phelps K, Didier ES, deHaro E, Marx PA, Traina-Dorge VL. 2012. Increased cellular immune responses and CD4+ T-cell proliferation correlate with reduced plasma viral load in SIV challenged recombinant simian varicella virus - simian immunodeficiency virus (rSVV-SIV) vaccinated rhesus macaques. Virology Journal 9: 160.

70. Das A, Veazey RS, Wang X, Lackner AA, Xu H, Pahar B. 2011. Simian immunodeficiency virus infection in rhesus macaques induces selective tissue specific B cell defects in double positive CD21+CD27+ memory B cells. Clinical Immunology 140: 223–228.

71. Das A, Xu H, Wang X, Yau CL, Veazey RS, Pahar B. 2011. Double-Positive CD21+CD27+ B Cells Are Highly Proliferating Memory Cells and Their Distribution Differs in Mucosal and Peripheral Tissues. PLoS ONE 6:e16524.

72. Schwan TG, Schrumpf ME, Hinnebusch BJ, Anderson DE, Konkel ME. 1996. GlpQ: an antigen for serological discrimination between relapsing fever and Lyme borreliosis. J Clin Microbiol 34: 2483–2492.

73. Lopez JE, Wilder HK, Hargrove R, Brooks CP, Peterson KE, Beare PA, Sturdevant DE, Nagarajan V, Raffel SJ, Schwan TG. 2013. Development of genetic system to inactivate a Borrelia turicatae surface protein selectively produced within the salivary glands of the arthropod vector. PLoS Negl Trop Dis 7:e2514.

